# Integrative Transcriptomic and Network Analysis Reveals Small Open Reading Frames as Regulators of DNA Methylation-Linked Pathogenicity and Adaptability in *Leptospira interrogans*

**DOI:** 10.64898/2025.12.08.692996

**Authors:** ChungYuen Khew, Nur Syafiqah Mohd Fowzi, Nurul Najihah Zaifulzaman, Sarahani Harun, Shairah Abdul Razak, Zeti-Azura Mohamed-Hussein, Norfarhan Mohd-Assaad

## Abstract

Small open reading frames (sORFs) are increasingly recognized as crucial regulators in bacterial gene expression, yet their biological roles remain largely unexplored in pathogenic species. Here, we investigated the genome-wide regulatory landscape of sORFs in Leptospira interrogans serovar Manilae strain UP-MMC-NIID-LP using RNA-seq–based transcriptomic profiling integrated with weighted gene co-expression network analysis (WGCNA) following targeted disruption of *lomA*, a gene mediating 4-methylcytosine (4mC) DNA modification. Loss of 4mC was associated with broad transcriptional dysregulation and phenotypic impairments, including reduced motility, adhesion, and virulence. Analysis of 363 predicted sORFs identified 39 with significant differential expression (FDR < 0.05, |log₂FC| ≥ 1) across wild-type, mutant, and complemented strains. Gene co-expression networks constructed using WGCNA and interrogated with Cytoscape tools (MCODE, CytoHubba, ClueGO) revealed ten differentially expressed sORFs (FDR-adjusted p < 0.05); eight upregulated and two downregulated, across the three pairwise comparisons of wild-type, mutant, and complemented strains. These sORFs were enriched in pathways related to flagellar assembly, DNA recombination, and transcriptional regulation, which are core processes supporting genome stability and adaptive stress responses. Several previously uncharacterized sORFs occupied hub-like positions within co-expression modules, highlighting their integrative roles in metabolic and regulatory networks. To our knowledge, this represents the first genome-wide integration of methylation-driven sORF regulation in Leptospira, revealing small proteins as central mediators that link epigenetic control with bacterial pathogenicity and adaptability. These findings provide a foundation for future antimicrobial and synthetic biology strategies targeting sORF-mediated regulation.

**Importance:** Small open reading frames (sORFs) are emerging as important regulators of bacterial gene expression, yet their roles in pathogenic species remain largely unexplored. This study provides the first genome-wide framework linking methylation-driven sORF regulation to virulence and adaptive processes in *Leptospira interrogans*. We identify previously uncharacterized sORFs occupying central positions in regulatory networks, connecting epigenetic control to motility, adhesion, and stress adaptation. Understanding these mechanisms in a neglected tropical disease pathogen has implications for improving public health and informs future antimicrobial and synthetic biology strategies.

## Introduction

Small open reading frames (sORFs) are short coding sequences, typically less than 300 nucleotides (nt) in length that encode proteins smaller than 100 amino acids (Leong et al., 2022). Although often overlooked due to their size, non-canonical start codons (CUG, GUG, UUG), and overlapping positions with other coding regions, sORFs are increasingly recognized as important regulators in both prokaryotic and eukaryotic systems (Lu et al., 2023). They occur in diverse genomic regions, including 5′ and 3′ untranslated regions (UTRs), intergenic spaces, and non-coding RNA regions (Jain et al., 2023). Advances in next-generation sequencing (NGS), ribosome profiling, and computational pipelines have enabled systematic detection of these elements, and functional studies in prokaryotes have revealed their involvement in stress adaptation and metabolic regulation. Examples include MgrB in Mg²⁺ ion uptake, YobF in heat stress adaptation, and PmrR in lipopolysaccharide (LPS) defense modification (Garai & Blanc-Potard, 2020; Khitun et al., 2019). sORF-encoded peptides can act through several mechanisms, such as stabilizing membrane complexes, modulating the activity of larger proteins, or functioning as signaling switches that fine-tune transcriptional or metabolic responses (Dong et al., 2023; Gray et al., 2022; Simoens et al., 2023). These micropeptides often interact transiently with protein complexes or membranes, exerting disproportionate regulatory influence despite their small size.

Building on these biological advances, recent methodological innovations have greatly expanded the capacity to systematically detect and validate sORFs across diverse organisms. Interest in sORFs has increased with evidence of their regulatory capacity and tractability as molecular agents. Recent studies demonstrated that small proteins are accessible to modern experimental and therapeutic tools (Perdikopanis et al., 2024). High-throughput discovery is driven by ribosome profiling (Ribo-Seq), which detects active translation events via three-nucleotide periodicity, and specialized mass spectrometry-based (MS) proteomics that directly identify micropeptide products (Kute et al., 2022). Following discovery, targeted methods employing antisense oligonucleotides (ASOs) and RNA interference (RNAi) strategies can be used to suppress sORF translation through mechanisms like steric hindrance or mRNA degradation for functional validation (Tsai et al., 2023; Xiao et al., 2025). In parallel, CRISPR- or phage-mediated genome editing systems offer precision tools to selectively disable or modify sORF loci in bacterial pathogens (Sun et al., 2024; Valdivia-Francia & Sendoel, 2024). Comprehensive Ribo-Seq and MS proteogenomics validation remain essential to confirm expression, structure, and accessibility of sORF products before functional interpretation (Zhang et al., 2022). Equally important, computational prediction pipelines enable rapid, unbiased genome-wide identification of candidate sORFs, providing a scalable foundation for mass discovery that guides and prioritizes downstream experimental validation. Collectively, these advances highlight the translational potential of the sORF candidates explored in this study.

Despite this expanding toolkit, the role of sORFs in bacterial pathogens with epigenetically regulated virulence remains underexplored. In *Leptospira interrogans*, a pathogenic spirochete responsible for leptospirosis, the small-protein landscape remains largely uncharacterized and its influence of epigenetic regulators such as *lomA* on their expression is unknown. The DNA methyltransferase *lomA* plays a central role in transcriptional regulation, growth, and virulence within this organism (Gaultney et al., 2020). Given these knowledge gaps, we asked whether the disruption of *lomA* alters sORF expression patterns and whether these sORFs map onto co-expression network modules associated with previously documented phenotypic in the *lomA* mutant, including disrupted motility, metabolic shifts, and impaired stress adaptation. *L. interrogans* provides a valuable model for investigating sORF-mediated regulation because it combines a moderately compact genome with a complex dual lifestyle, surviving in environmental reservoirs while infecting mammalian hosts. Its virulence is epigenetically regulated through DNA methylation, implicating *lomA* as a regulatory hub (Gaultney et al., 2020). Leptospirosis is a widespread zoonotic disease with more than one million cases and approximately 60,000 deaths annually (Khew et al., 2023). In Malaysia, it ranks as third leading infection cause of death after dengue and malaria (Philip & Ahmed, 2023). Transcriptomic comparisons between pathogenic vs saprophytic *Leptospira* species have revealed shifts in motility, stress response and host-adaptation programs, yet these studies excluded the small-protein coding landscape (Davignon et al., 2024; Giraud-Gatineau et al., 2024).

To investigate this potential regulatory link between *lomA* and the sORF landscape, we reanalyzed RNA-Seq data (GEO: GSE138917) encompassing wild-type, *lomA* mutant, and complemented *L. interrogans* strains, enabling causal inference of *lomA*-linked expression changes rather than simple correlation. Notably, *lomA* is conserved across pathogenic and intermediate *Leptospira* species but absent from saprophytic species, implicating its virulence regulation (Gaultney et al., 2020). Disruption of *lomA* leads to global transcriptional dysregulation, reduced growth, impaired adhesion, and loss of virulence in infection models (Gaultney et al., 2020). We integrated RNA-Seq transcriptomic profiling with weighted gene co-expression network analysis (WGCNA) to identify co-expressed gene modules and candidate hub-like sORFs. WGCNA is well suited here because its groups genes with correlated expression patterns into modules and allows us to detect hub-genes that coordinate pathways (Heidarzadehpilehrood et al., 2023; Langfelder & Horvath, 2008).

This study presents the first genome-wide integration of methylation-driven sORF regulation in *L. interrogans*. Our systems-level approach bridges *lomA*-dependent DNA methylation with small-protein expression, revealing how sORFs contribute to bacterial adaptation and virulence. Combining RNA-Seq transcriptomic profiling with WGCNA and functional enrichment uncovered sORFs embedded within gene network modules linked to motility, nucleotide metabolism, and virulence regulation. Extending this analysis to sORFs overlapping with small RNAs broadened the search beyond standard annotation filters, allowing the detection of functional coding elements often missed in conventional analyses. This work fills a critical gap in leptospiral transcriptomics by demonstrating that both annotated and uncharacterized sORFs play integral roles within the regulatory networks disrupted by *lomA* inactivation. Our findings highlight the potential of sORFs to act as regulatory nodes in host adaptation and pathogenesis, offering new perspectives for therapeutic targeting and genome annotation refinement.

## Methods

### Small ORFs prediction and annotation

To identify small open reading frames (sORFs) in *Leptospira interrogans*, we utilized RNA-Seq datasets (GSE138917) from NCBI’s GEO database, which included samples from wild-type, complemented strains, and *lomA* mutant. The *L. interrogans* serovar Manilae reference genome (NCBI Accession: GCF_001047635.1), sequenced via PacBio technology and annotated following Gaultney et al. (2020), was used as the genomic framework. Initial read quality was assessed with FastQC v1.17 (Andrews, 2010), complemented by MultiQC summary reports to verify dataset integrity. Adapter trimming and quality filtering were performed using Trimmomatic v0.39 (Ewels et al., 2016) in single-end (SE) mode with the following parameters: (LEADING:3, TRAILING:3, SLIDINGWINDOW:4:15, MINLEN:30).

Cleaned reads were aligned to the reference genome via the READemption v2.0.4 (Förstner et al., 2014) pipeline, employing default parameters for single-end reads optimized for bacterial transcriptome analysis. Subsequent transcript assembly, sRNA, and sORF prediction were performed using ANNOgesic v1.0.22 (Yu et al., 2018) with default parameters, leveraging strand-specific coverage data for each replicate. The “-print_all_combination” option was applied during sORF detection to comprehensively evaluate candidate ORFs. Non-coding RNA candidates were systematically excluded to refine the sORF candidate pool, facilitating focused functional annotation. Annotation integration utilized BLAST2GO (Conesa & Götz, 2008) and InterProScan (Jones et al., 2014) to assign gene ontology terms and domain-based functions, underpinning downstream biological interpretation.

### Transcriptome analyses

Trimmed single-end reads were aligned to *L. interrogans* reference genome using STAR v2.7.11a (Dobin et al., 2013), employing strand-specific settings and genome indices built from a merged annotation combining NCBI and ANNOgesic output. Post-alignment BAM files were sorted and indexed with SAMtools v1.0 (Li et al., 2009), facilitating compatibility with quantification tools. Gene-level expression quantification was performed using HTSeq-Count (Anders et al., 2015) in single-end mode, applying the union counting method to maximize assignment precision. Differential expression analyses were conducted using the edgeR v4.2.0 package (Robinson et al., 2009), implementing normalization and dispersion estimation following best practices. Comparisons among wild-type, complement, and mutant strains revealed transcriptional changes, with significance thresholds set at a false discovery rate (FDR) < 0.05 and |log₂ fold change| ≥ 1. Visualization of differentially expressed genes employed volcano plots generated with ggplot2 in R, providing clear depictions of transcriptional dynamics.

### Construction and functional analysis of sORF-associated co-expression networks

Functional inference for sORFs was performed by integrating gene annotation data with differential expression profiles. Genes exhibiting statistically significant regulation (adjusted *p* < 0.05, |log₂FC| ≥ threshold) were prioritized for network-based analysis to identify potential roles in virulence, metabolic adaptation, and cellular regulation in *L. interrogans*. Weighted gene co-expression network analysis (WGCNA) was conducted using WGCNA package in R with the “*goodSamplesGenes*” function to remove low-quality entries (Langfelder & Horvath, 2008). A soft-thresholding power of 12 was selected to approximate a scale-free topology, and the resulting adjacency matrix was transformed into a topological overlap matrix (TOM) for hierarchical clustering. Modules of co-expressed genes were defined using dynamic tree cutting, and module eigengenes were calculated to summarize expression profiles (Langfelder & Horvath, 2014).

Network visualization and functional enrichment were performed in Cytoscape v3.10.2 (Shannon et al., 2003). Weighted gene co-expression network modules were first exported as gene–gene interaction tables, and these were used as input for downstream network analysis. Highly interconnected clusters were identified using the MCODE plugin with the following parameters: degree cutoff = 2, node score cutoff = 0.2, K-core = 2, and maximum depth = 100 (Bader & Hogue, 2003). Putative hub-like genes were ranked using the CytoHubba plugin, taking the full module-derived networks as input and applying the Maximal Clique Centrality (MCC) algorithm, which prioritizes nodes by clique connectivity (Chin et al., 2014). In this study, the sORFs with the top 10 MCC values were considered as hub-like genes. Functional annotation and enrichment were performed with the ClueGO plugin (Bindea et al., 2009), integrating Gene Ontology terms (biological process, molecular function, cellular component, immune system) and KEGG pathways. Default statistical settings were applied, including a two-sided hypergeometric test, Benjamini–Hochberg correction for multiple testing (*p* < 0.05), Kappa score threshold = 0.4, minimum three genes per term, and a minimum of 4% representation per term. The latest *Leptospira interrogans* serovar Manilae strain UP-MMC-NIID-LP genome annotation was used to ensure functional accuracy.

## Results

### Identification and Functional Annotation of sORFs in *Leptospira interrogans*

Using the ANNOgesic pipeline (Yu et al., 2018), a comprehensive genome scan identified a total of 1,253 putative sORFs from the integrated RNA-Seq data (Table 1). Of these, 363 high-confidence sORF candidates were retained based on stringent criteria prioritizing sequence length (<300 nt or 100 amino acids), presence of ribosomal binding sites (RBS) and overlap with sRNA. Ensuring these sORFs encode peptides rather than being noncoding fragments, we required the presence of Shine-Dalgarno sequences (unique to prokaryotes), start and stop codons within the non-coding expressed regions, following established bacterial translation initiation models (Fuchs & Engelmann, 2023; Giess et al., 2017). Since recent reports indicate sRNAs contain translatable sORFs (Aoyama et al., 2022; George et al., 2024), we included overlapping candidates, adding 122 candidate sequences to the high confidence set (241 non-overlapping versus 363 total). Across the 363 predicted sORFs, majority exhibited very short nucleotide lengths, with 307 sequences (85%) measuring ≤100 nt. An additional 51 sORFs (14%) fell within the 101–180 nt range, and only 5 sORFs (1%) within the range of 181–300 nt. The strong representation of short sORFs is consistent with previously reported bacterial sORF length profiles.

**Table 1.** Summary of predicted and filtered small open reading frames (sORFs) across the complete genome. A total of 1,253 putative sORFs were identified genome-wide, stratified by genomic location (chromosomes and plasmid) and genomic context (intergenic or antisense). *\*Percentage indicates the proportion of best sORFs that do not overlap predicted sRNAs, calculated as (non-overlapping best sORFs / total best sORFs) × %*.

| Region / Context | Total sORFs | Total sORFs (No sRNA Overlap) | Total sORFs (with RBS) | Total sORFs (with RBS & No sRNA Overlap) | Best sORFs (No sRNA Overlap) * | Best sORFs (with RBS & sRNA overlap) |
| --- | --- | --- | --- | --- | --- | --- |
| **Whole Genome** | 1,253 | 371 | 363 | 257 | 241 (66.4%) | 363 |
| Intergenic | 404 | 95 | 99 | 70 | 63 (63.6%) | 99 |
| Antisense | 849 | 276 | 264 | 187 | 178 (67.4%) | 264 |
| **Chromosome 1** | 1,102 | 325 | 320 | 227 | 207 (64.7%) | 320 |
| Intergenic | 350 | 83 | 87 | 62 | 56 (64.4%) | 87 |
| Antisense | 752 | 242 | 233 | 165 | 151 (64.8%) | 233 |
| **Chromosome 2** | 125 | 41 | 38 | 13 | 29 (76.3%) | 38 |
| Intergenic | 37 | 10 | 9 | 3 | 4 (44.4%) | 9 |
| Antisense | 88 | 31 | 29 | 10 | 25 (86.2%) | 29 |
| **Plasmid** | 26 | 5 | 5 | 2 | 5 (100%) | 5 |
| Intergenic | 17 | 2 | 3 | 0 | 3 (100%) | 3 |
| Antisense | 9 | 3 | 2 | 2 | 2 (100%) | 2 |

Sequence-based annotation using BLAST2GO against the NCBI non-redundant (nr) database returned significant hits for 101 (27.8%) candidates, while 20 of the 363 sORFs (5.5%) were associated with InterProScan domain hits, many of which corresponded to predicted transmembrane segments or signal peptide features (Table 2). InterProScan v5, which integrates PHOBIUS, TMHMM, HMMER-based profiles and multiple domain libraries, was used for domain and topology prediction. These findings hint at a possible enrichment of membrane-associated or regulatory functions among the most confidently predicted sORFs, warranting further experimental validation. An independent analysis using the InterProScan webserver, focusing on TMHMM analysis further supported this, identifying four sORFs encoding proteins with single alpha-helical transmembrane segments, characteristic of inner membrane proteins in Gram-negative bacteria (Garai & Blanc-Potard, 2020). Notably, single alpha-helical transmembrane segments are often associated with regulatory or signaling roles rather than classical transport, as such single-pass proteins frequently function as modulators of protein complexes or signal transducers (Yadavalli & Yuan, 2022).

**Table 2.**
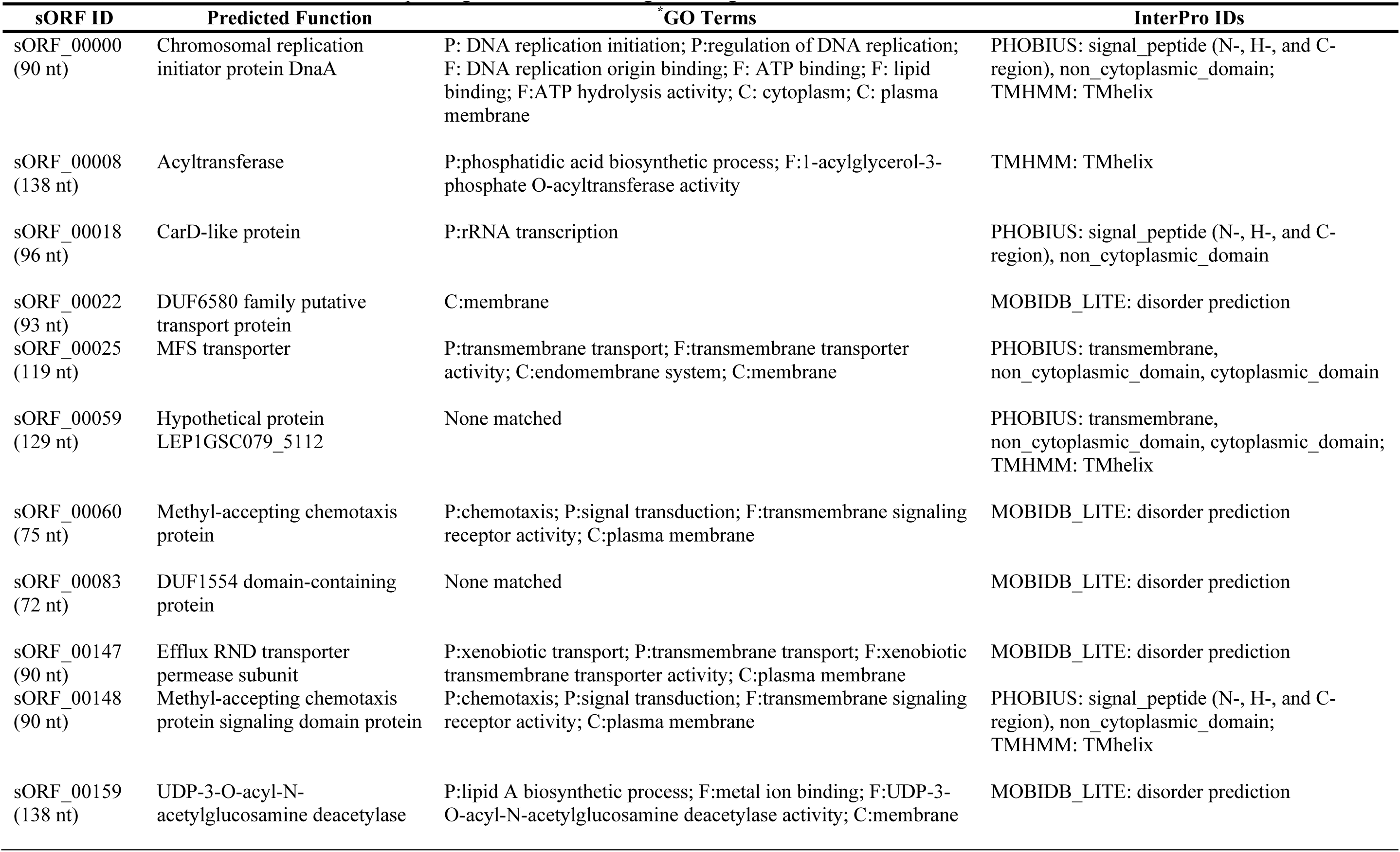
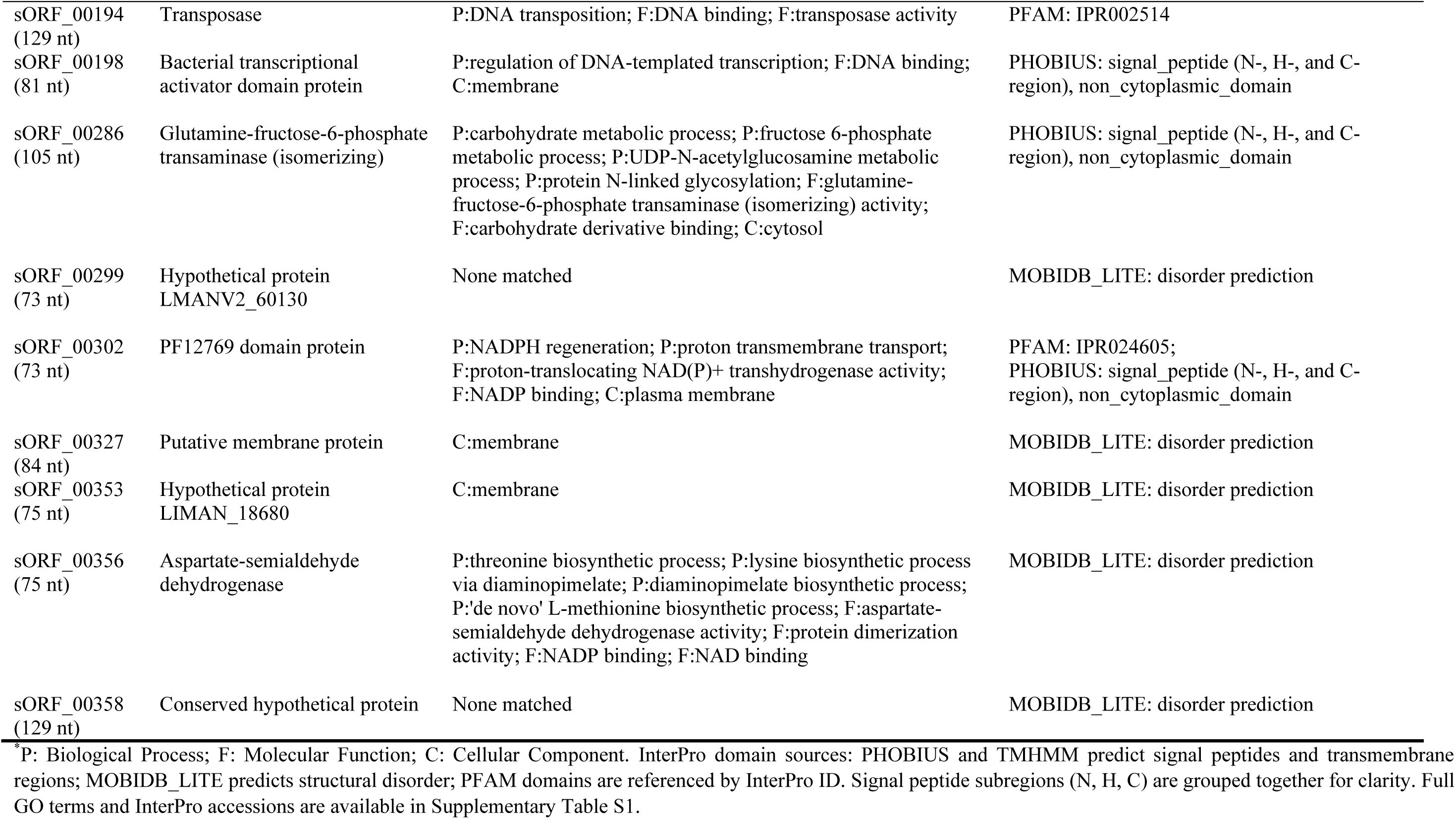
Functional annotation of 20 sORFs associated with InterProScan-predicted domains. These 20 sORFs (5.5% of the high-confidence set) returned InterProScan domain identifiers, with most annotations corresponding to transmembrane regions or signal transduction-associated features.

| sORF ID | Predicted Function | *GO Terms | InterPro IDs |
| --- | --- | --- | --- |
| sORF_00000<br>(90 nt) | Chromosomal replication initiator protein DnaA | P: DNA replication initiation; P:regulation of DNA replication; F: DNA replication origin binding; F: ATP binding; F: lipid binding; F:ATP hydrolysis activity; C: cytoplasm; C: plasma membrane | PHOBIUS: signal_peptide (N-, H-, and C-region), non_cytoplasmic_domain; TMHMM: TMhelix |
| sORF_00008<br>(138 nt) | Acyltransferase | P:phosphatidic acid biosynthetic process; F:1-acylglycerol-3-phosphate O-acyltransferase activity | TMHMM: TMhelix |
| sORF_00018<br>(96 nt) | CarD-like protein | P:rRNA transcription | PHOBIUS: signal_peptide (N-, H-, and C-region), non_cytoplasmic_domain |
| sORF_00022<br>(93 nt) | DUF6580 family putative transport protein | C:membrane | MOBIDB_LITE: disorder prediction |
| sORF_00025<br>(119 nt) | MFS transporter | P:transmembrane transport; F:transmembrane transporter activity; C:endomembrane system; C:membrane | PHOBIUS: transmembrane, non_cytoplasmic_domain, cytoplasmic_domain |
| sORF_00059<br>(129 nt) | Hypothetical protein LEP1GSC079_5112 | None matched | PHOBIUS: transmembrane, non_cytoplasmic_domain, cytoplasmic_domain; TMHMM: TMhelix |
| sORF_00060<br>(75 nt) | Methyl-accepting chemotaxis protein | P:chemotaxis; P:signal transduction; F:transmembrane signaling receptor activity; C:plasma membrane | MOBIDB_LITE: disorder prediction |
| sORF_00083<br>(72 nt) | DUF1554 domain-containing protein | None matched | MOBIDB_LITE: disorder prediction |
| sORF_00147<br>(90 nt) | Efflux RND transporter permease subunit | P:xenobiotic transport; P:transmembrane transport; F:xenobiotic transmembrane transporter activity; C:plasma membrane | MOBIDB_LITE: disorder prediction |
| sORF_00148<br>(90 nt) | Methyl-accepting chemotaxis protein signaling domain protein | P:chemotaxis; P:signal transduction; F:transmembrane signaling receptor activity; C:plasma membrane | PHOBIUS: signal_peptide (N-, H-, and C-region), non_cytoplasmic_domain; TMHMM: TMhelix |
| sORF_00159<br>(138 nt) | UDP-3-O-acyl-N-acetylglucosamine deacetylase | P:lipid A biosynthetic process; F:metal ion binding; F:UDP-3-O-acyl-N-acetylglucosamine deacetylase activity; C:membrane | MOBIDB_LITE: disorder prediction |
| sORF_00194<br>(129 nt) | Transposase | P:DNA transposition; F:DNA binding; F:transposase activity | PFAM: IPR002514 |
| sORF_00198<br>(81 nt) | Bacterial transcriptional activator domain protein | P:regulation of DNA-templated transcription; F:DNA binding; C:membrane | PHOBIUS: signal_peptide (N-, H-, and C-region), non_cytoplasmic_domain |
| sORF_00286<br>(105 nt) | Glutamine-fructose-6-phosphate transaminase (isomerizing) | P:carbohydrate metabolic process; P:fructose 6-phosphate metabolic process; P:UDP-N-acetylglucosamine metabolic process; P:protein N-linked glycosylation; F:glutamine-fructose-6-phosphate transaminase (isomerizing) activity; F:carbohydrate derivative binding; C:cytosol | PHOBIUS: signal_peptide (N-, H-, and C-region), non_cytoplasmic_domain |
| sORF_00299<br>(73 nt) | Hypothetical protein LMANV2_60130 | None matched | MOBIDB_LITE: disorder prediction |
| sORF_00302<br>(73 nt) | PF12769 domain protein | P:NADPH regeneration; P:proton transmembrane transport; F:proton-translocating NAD(P)+ transhydrogenase activity; F:NADP binding; C:plasma membrane | PFAM: IPR024605; PHOBIUS: signal_peptide (N-, H-, and C-region), non_cytoplasmic_domain |
| sORF_00327<br>(84 nt) | Putative membrane protein | C:membrane | MOBIDB_LITE: disorder prediction |
| sORF_00353<br>(75 nt) | Hypothetical protein LIMAN_18680 | C:membrane | MOBIDB_LITE: disorder prediction |
| sORF_00356<br>(75 nt) | Aspartate-semialdehyde dehydrogenase | P:threonine biosynthetic process; P:lysine biosynthetic process via diaminopimelate; P:diaminopimelate biosynthetic process; P:'de novo' L-methionine biosynthetic process; F:aspartate-semialdehyde dehydrogenase activity; F:protein dimerization activity; F:NADP binding; F:NAD binding | MOBIDB_LITE: disorder prediction |
| sORF_00358<br>(129 nt) | Conserved hypothetical protein | None matched | MOBIDB_LITE: disorder prediction |

\*P: Biological Process; F: Molecular Function; C: Cellular Component. InterPro domain sources: PHOBIUS and TMHMM predict signal peptides and transmembrane regions; MOBIDB\_LITE predicts structural disorder; PFAM domains are referenced by InterPro ID. Signal peptide subregions (N, H, C) are grouped together for clarity. Full GO terms and InterPro accessions are available in Supplementary Table S1.

A large fraction (262/363; 72.2%) lacked detectable homology, consistent with the taxonomically restricted and rapidly evolving nature of bacterial small proteins, which often lack conserved domains yet can retain regulatory function (Leong et al., 2022; Yadavalli & Yuan, 2022). It suggests these sORFs may be taxonomically restricted or represent lineage-specific microproteins (Simoens et al., 2023). The full results for all 363 sORF candidates predicted by ANNOgesic are provided in **Supplementary Table S1**. In addition to the absence of homology, many of these “hypothetical” sORFs show predicted transmembrane or signal-peptide features, which may provide functional plausibility that small membrane-associated proteins frequently act as signaling adaptors or local regulators of larger protein complexes (Yadavalli & Yuan, 2022). Detailed set of BLAST and InterProScan annotations corresponding to these candidates is available at **Supplementary Table S2**.

### Expression Profile of sORFs in *L. interrogans*

To assess the transcriptional impact of *lomA* disruption, RNA-Seq data from *L. interrogans* serovar Manilae strain UP-MMC-NIID-LP (GEO accession GSE138917), originally generated by Gaultney et al. (2020) were reanalyzed. This dataset comprises three transcriptomes of wild-type, *lomA* mutant, and complemented strains, providing a robust framework to evaluate differential gene expression of methylation-linked changes. Reads were mapped to the reference genome, and transcript abundances were quantified using standard pipelines prior to downstream statistical analyses as detailed in Methodology.

Differential expression profiling across the three conditions identified 157 non-redundant differentially expressed genes (DEGs), following false discovery rate (FDR) < 0.05 and |log₂ fold change| ≥ 1 (Table 3). This sum accounts for overlapping genes across pairwise comparisons, ensuring each differentially expressed gene is counted only once. Contrasting expression shifts were observed between the *lomA* mutant and wild-type pairwise, highlighting the regulatory impact of *lomA* on the global gene transcription of *L. interrogans* consistent with Gaultney et al. (2021). Among these, 39 were sORFs, including both annotated and newly identified (unknown) sequences. Of the 39, 30 (approximately 77%) were upregulated in the *lomA* mutant relative to the wild-type strain, indicating that loss of *lomA*-mediated DNA methylation broadly de-represses sORF expression.

**Table 3.**
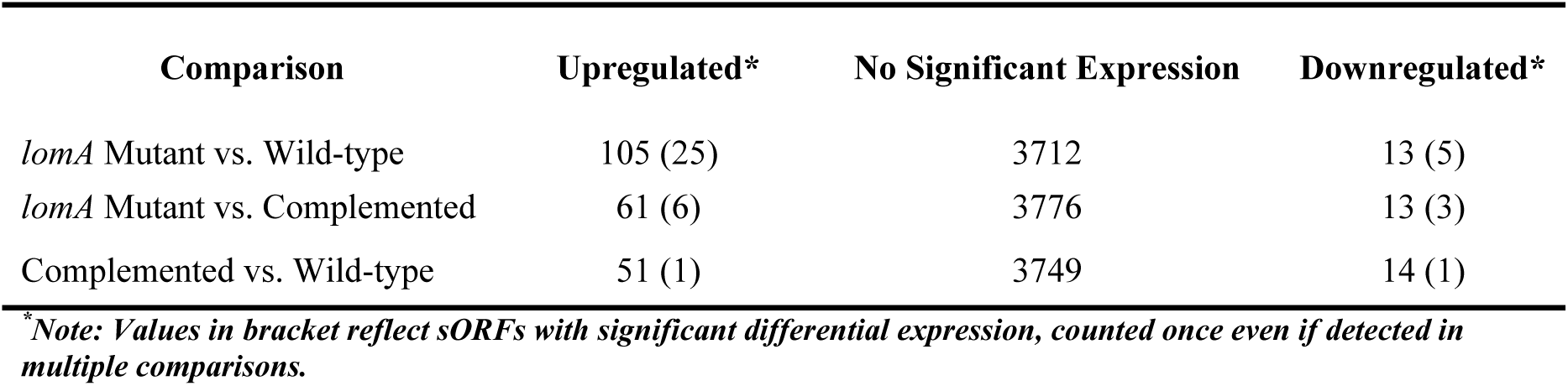
Expression changes in *lomA* mutant compared to wild-type and complemented strain. Comparative expression data demonstrate the extent of transcriptional changes induced by the *lomA* mutation, highlighting both upregulation and downregulation of genes relative to control strains and revealing the regulatory impact of *lomA* in *L. interrogans*.

Notably, sORF_00332 exhibited one of the strongest transcriptional inductions (>32-fold) following *lomA* disruption (mutant-wildtype pairwise), suggesting sensitivity to methylation-dependent regulation and potential biological significance in addition to statistical relevance. Although BLAST2GO provided no functional annotation, it’s robust upregulation in a context where *lomA* disruption reduces virulence and motility (Gaultney et al., 2020), positions sORF_00332 as a prime candidate for methylation-sensitive regulatory or effector roles, warranting prioritization for experimental validation through tagged expression or knockout assays. Other sORFs, such as sORF_00010 (CoA ligase-related), sORF_00166 (ATP-binding), and sORF_00331 (putative adenosylhomocysteinase), exhibited substantial expression shifts, indicative of functions in catalysis, transport, and stress response.

Volcano plots visualized differential expression across each pairwise conditions (Fig. 1). The mutant-wildtype contrast revealed the greatest transcriptional disruption, with 105 upregulated and 13 downregulated genes, including 30 sORFs (25 upregulated and five downregulated). This suggests that sORFs were disproportionately affected by the mutation compared to the wildtype strain. In the mutant-complemented comparison, 61 genes were upregulated and 13 downregulated, including nine sORFs were differentially expressed (six upregulated and three downregulated), showing partial restoration of expression levels upon complementation. The complemented-wildtype pairwise showed 51 genes were significantly upregulated and 14 downregulated, but only two sORFs were differentially expressed (one upregulated and downregulated respectively), suggesting greater recovery of transcriptional balance. A full list of differentially expressed sORFs across all contrasts is provided in **Supplementary Table S3-S7**.

**Fig. 1.**
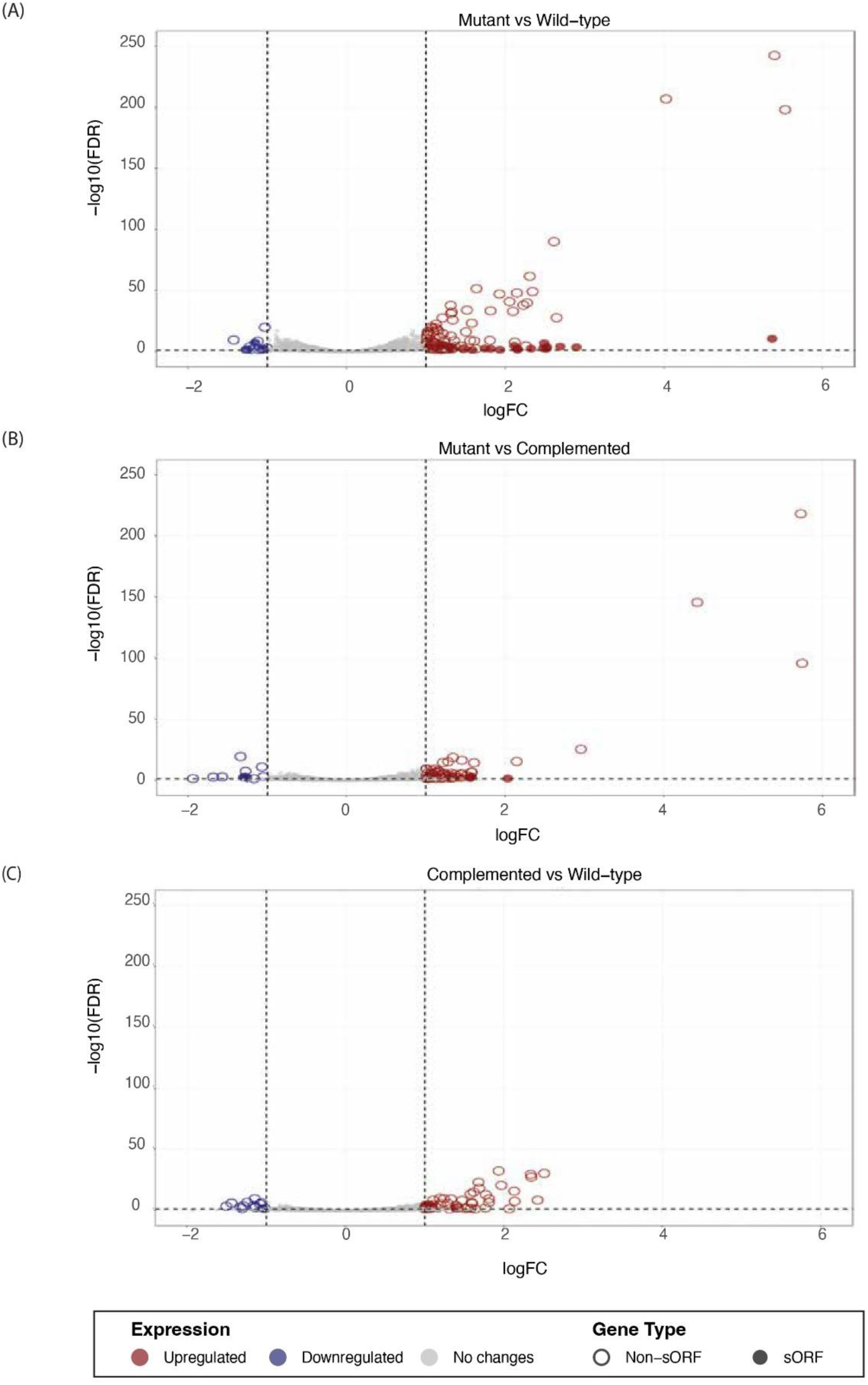
Differential expression profiles across wild-type, mutant, and complemented strains based on FDR < 0.05 and |log₂FC| ≥ 1 thresholds. Volcano plots display differentially expressed genes (DEGs) across three pairwise comparisons: (A) mutant vs. wild-type, (B) mutant vs. complemented, and (C) complemented vs. wild-type. The plots illustrate the distinct transcriptional impact of the mutation and its partial restoration upon complementation.

### Weighted Gene Co-expression Network Analysis Reveals sORF-Centric Regulatory Modules in Bacterial Mutants

Using the WGCNA R package, a topological overlap matrix (TOM) was computed to quantify gene connectivity, and 64 co-expression modules were identified. Modules contained minimum of 30 highly correlated genes and were assigned distinct color labels. The ‘grey’ module, lacking detectable gene interactions, and the ‘mediumorchid’ module, which did not meet default ClueGO parameters, were excluded from further analysis. Of the remaining 62 modules, 41 passed ClueGO enrichment under default parameters, which include a two-sided hypergeometric test with Benjamini-Hochberg correction (*p* < 0.05), a Kappa score threshold of 0.4, and a minimum of three genes per term with ≥4% representation. Twenty-five modules contained one or more sORFs, and six were tightly associated with *lomA*-dependent transcriptional changes.

Integration of WGCNA with CytoHubba Maximal Clique Centrality (MCC) ranking revealed multiple sORFs occupying central module positions. Hub-like sORFs were defined based on network topology using the CytoHubba plugin in Cytoscape, specifically, the top 10 nodes by MCC score within each module. As this classification is derived from co-expression topology rather than biochemical validation, these sORFs are referred to as hub-like elements throughout. Upregulated sORFs were detected in the ‘blue’, ‘plum1’, ‘red’, ‘skyblue’, and ‘turquoise’ modules, whereas downregulation was observed in the ‘purple’ module (Table 4). Their central positioning in each module supports the view that small proteins may act as regulatory nodes coordinating adaptive responses related to membrane remodeling, chemotaxis, and nucleotide metabolism. Such putative functions are consistent with prior observations that small membrane-associated proteins can influence larger complexes and signaling cascades though these relationships in *Leptospira* remain to be directly demonstrated (Burton et al., 2024; Hemm et al., 2020).

**Table 4.**
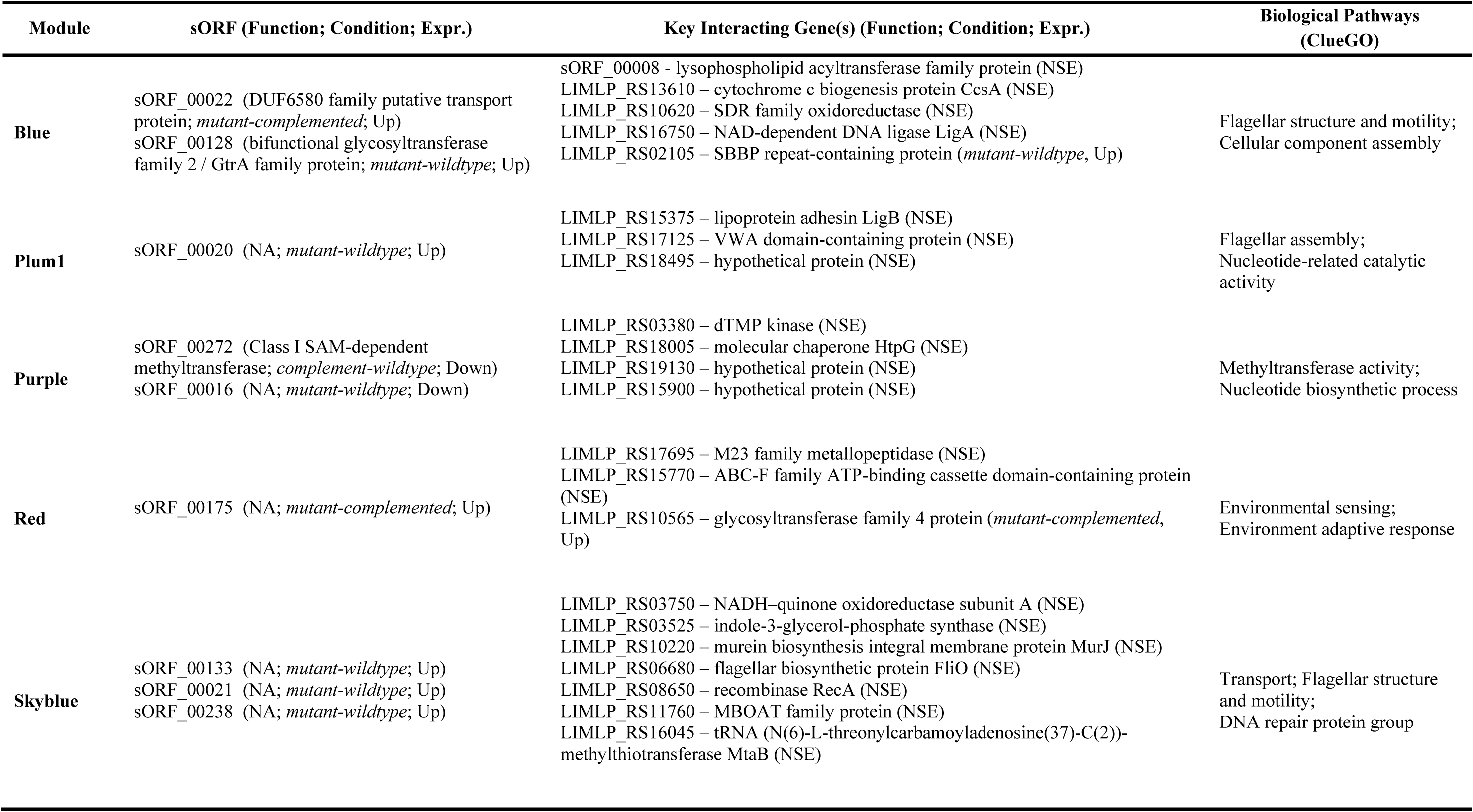
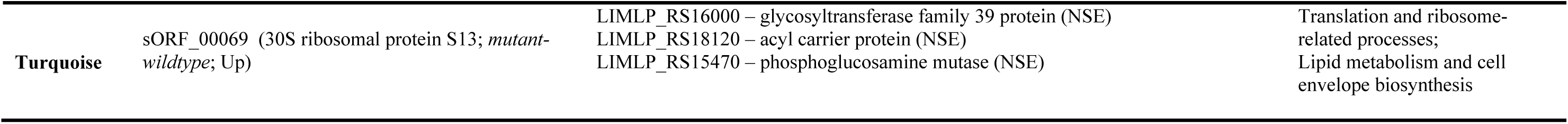
Summary of Modules with Differentially-Expressed sORF, Key Interacting Genes, and Associated Biological Pathways. Summary of identified sORFs from WGCNA-Cytoscape analysis, their key interacting genes, and ClueGO-derived biological pathway associations. Functional annotation, experimental condition, and expression pattern (Up, Down, NSE = no significant expression) are shown for each gene.

While functional inference in this study is based on GO-term enrichment and sequence homology, direct biochemical validation will be required to confirm the regulatory roles of these hub sORFs. Nevertheless, the predictive use of WGCNA in bacterial transcriptomics has been well established for identifying co-regulated clusters potentially involved in stress responses and surface architecture remodeling adaptation (Heidarzadehpilehrood et al., 2023; Lu et al., 2021). Collectively, these findings identify candidate sORFs that may participate in methylation-sensitive regulatory circuits, but confirmation of such integrative roles will require functional assays. Detailed molecular interactions for each differentially expressed sORF, together with their strongest weighted interactors and enriched biological processes, are summarized in Table 4, while corresponding ClueGO functional networks are shown in Fig. 2A–F.

**Fig. 2.**
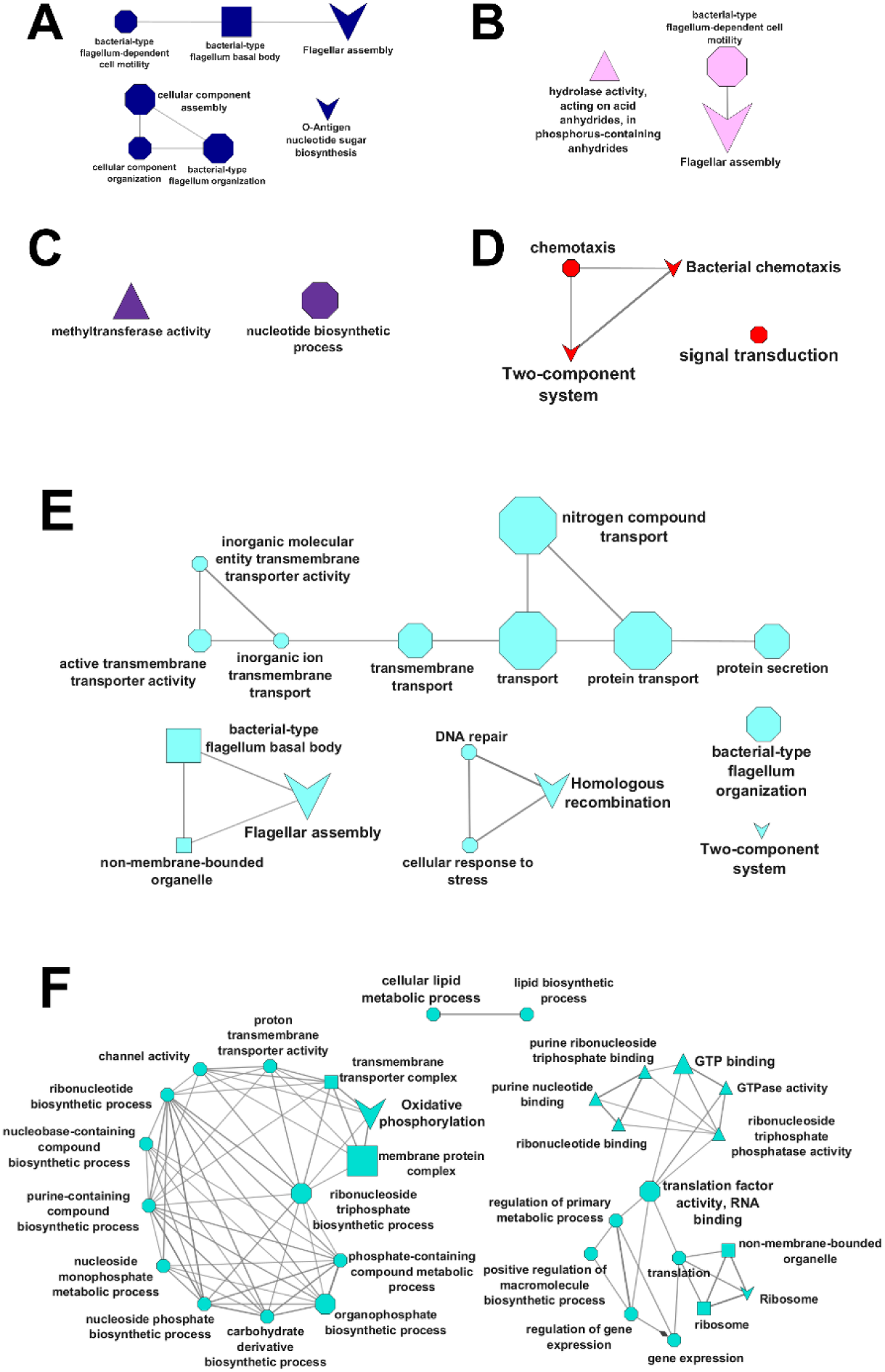
Functional association networks of representative WGCNA modules enriched for differentially expressed sORFs. **(A)** *Blue module*: Upregulated sORF_00022 and sORF_00128 enriched for flagellar assembly and cell organization, with key interactions involving sORF_00008 and LIMLP_RS16750, indicating integration between motility and structural components. **(B)** *Plum1 module*: Upregulated sORF_00020 associated with flagellar cap formation, showing interaction with LIMLP_RS15375 (LigB), suggesting a link between flagellar morphogenesis and host adhesion. **(C)** *Purple module*: Downregulated sORF_00272 and sORF_00016 cluster within methyltransferase-related processes, interacting with LIMLP_RS03380 (dTMP kinase), implying potential roles in DNA modification or nucleotide metabolism. **(D)** *Red module*: Upregulated sORF_00175 is associated with chemotaxis and two-component signaling pathways, interacting with cheR and M23 metallopeptidase, supporting a potential role in signal-mediated motility control. **(E)** *Skyblue module*: Upregulated sORFs sORF_00133, sORF_00021, and sORF_00238 are involved in flagellar assembly and nitrogen transport, interacting with LIMLP_RS03750 and FliO, pointing to coordinated regulation of motility and nutrient flux. **(F)** *Turquoise module*: Upregulation of sORF_00069 linked to oxidative phosphorylation, with high-confidence interaction with LIMLP_RS16000 (glycosyltransferase), suggesting adaptive energetic remodeling.

### Functional Characterization of Key sORF-Linked Modules

In the ‘blue’ module (Fig. 2A), sORF_00022 was upregulated in the mutant-complemented comparison, suggesting restoration or activation of membrane transport pathways likely linked to CcsA-mediated cytochrome c biogenesis. Upregulation of SBBP repeat-containing protein (LIMLP_RS02105) in the mutant–wildtype contrast, indicative of increased expression of surface or structural proteins potentially influencing motility structures (Cinar et al., 2024). Within this module, sORF_00128, annotated as a bifunctional glycosyltransferase of the GtrA family, was found to interact with three proteins: lysophospholipid acyltransferase, DNA ligase (LigA), and an SBBP repeat-containing protein, being the only one upregulated in the mutant–wildtype comparison. Notably, sORF_00008 (lysophospholipid acyltransferase) emerged listed among the top three strongest-weight interactors for sORF_00022 (Table 4), emerged as a shared interactor between these gene groups, suggesting a central role in membrane remodeling that bridges lipid modification, glycosylation, and flagellar assembly, where similar mechanisms were also reported in *E. coli* (Niu et al., 2024; Toyotake et al., 2020).

In the ‘plum1’ module (Fig. 2B), sORF_00020 was connected to adhesion-related proteins such as LigB and VWA-domain-containing proteins. While none of its direct interactors were classical flagellar proteins, the enrichment for flagellar assembly and ATP/GTP hydrolase activity suggested indirect coupling between adhesion and motility functions. sORF_00020 was upregulated in the mutant–wildtype comparison, indicating condition-specific activation potentially linked to energy-dependent surface complex assembly. The ‘purple’ module (Fig. 2C) contains two functionally linked sORFs: sORF_00272, a Class I SAM-dependent methyltransferase, and sORF_00016, unannotated but co-clustered with dTMP kinase (LIMLP_RS03380) and HtpG. The network topology indicated a mechanistic bridge between nucleotide biosynthesis and nucleic acid modification, with both sORFs downregulated in mutant–wildtype contrasts, suggesting reduced DNA synthesis and modification functions.

In the ‘red’ module (Fig. 2D), uncharacterized sORF_00175 interacted with M23 metallopeptidase, glycosyltransferase family 4 protein, and an ABC-F family protein. Co-upregulation of sORF_00175 and the glycosyltransferase in the mutant-complemented comparison pointed to coordinated regulation of cell envelope composition, a process linked to chemotaxis and two-component signal transduction enrichment in the module. The ‘skyblue’ module (Fig. 2E) included three unannotated sORFs, namely sORF_00133, sORF_00021, and sORF_00238, which collectively integrate transport energetics, motility, DNA repair, and translation adaptation. sORF_00133 was consistently upregulated in mutant–wildtype contrasts and interacted with nuoA and MurJ, suggesting modulation of membrane transport and cell wall biosynthesis. sORF_00021 linked motility (FliO) with genome stability (RecA) and lipid modification (MBOAT). sORF_00238 shared oxidative phosphorylation links with sORF_00133 but also engaged tRNA modification machinery.

In the ‘turquoise’ module (Fig. 2F), sORF_00069, encoding ribosomal protein S13, was upregulated in mutant–wildtype comparisons and interacted with enzymes for cell envelope biosynthesis such as glycosyltransferase family 39 protein, acyl carrier protein, and phosphoglucosamine mutase. The module was enriched for both translation and lipid/cell wall biosynthetic processes, indicating coordinated regulation of protein synthesis and envelope assembly.

Following the identification of co-expression modules and key interactors, we next prioritized the most central sORFs using network topology metrics to distinguish highly connected elements from peripheral nodes. Building on the module-level patterns described above, Maximal Clique Centrality (MCC) scoring in CytoHubba was used to rank all nodes by intramodular connectivity, providing a more quantitative measure of network influence. In this framework, the top 10 ranked genes within each module were defined as hub-like sORFs, representing the most topologically central candidates likely to coordinate information flow within the module. This refined criterion minimizes overinterpretation by focusing on the highest-confidence nodes supported by network metrics rather than differential expression alone.

### Hub-like sORFs Identified by MCC Ranking Reveal Central Roles in Motility and Metabolic Regulation

Hub gene analysis of the WGCNA modules highlighted a diverse set of sORFs occupying central network positions, underscoring their potential regulatory influence across distinct bacterial processes (Table 5). Across modules, hub-like sORFs displayed moderate connection strengths (r = 0.3–0.45). Hub-like classification was based on MCC scores computed in CytoHubba, with the top ten ranked nodes per module defined as hub-like elements.

**Table 5.** Hub gene characteristics from CytoHubba MCC analysis within WGCNA modules. Summary of top-ranked hub-like sORFs identified within each WGCNA module using the Maximal Clique Centrality (MCC) method in CytoHubba.

| Gene ID (Module) | Predicted Functional Annotation | MCC Rank | Module GO Enrichment (ClueGO) | No. of First Neighbors (Nodes) | Network Size (Nodes) |
| --- | --- | --- | --- | --- | --- |
| sORF_00008 (Blue) | Lysophospholipid acyltransferase | 7th | Flagellar assembly; O-antigen nucleotide sugar biosynthesis | 108 | 110 |
| sORF_00284 (Purple) | NA | 5th | Methyltransferase activity; Nucleotide biosynthetic process | 90 | 92 |
| sORF_00175 (Red) | NA | 8th | Two-component system; Bacterial chemotaxis | 99 | 100 |
| sORF_00203 (Green) | NA | 3rd | Cytoplasm/cytosol localization; Carboxylic acid metabolism; Lipid/terpenoid biosynthesis; tRNA modification (supporting) | 83 | 103 |

In the ‘blue’ module, top-ranked hub-like sORF_00008 (lysophospholipid acyltransferase), exhibited strong associations with flagellar assembly and O-antigen nucleotide sugar biosynthesis, indicating a potential involvement in both motility apparatus assembly and surface antigen modification based on shared enrichment patterns. In the ‘purple’ module, the top-ranked sORF_00284 showed enrichment for methyltransferase activity and nucleotide biosynthetic processes, suggesting a role in epigenetic or metabolic regulation, although this remains speculative. In the ‘red’ module, the putative hub-like sORF_00175 showed enrichment for two-component system and chemotaxis-related GO terms, suggesting a potential, though unverified, link between environmental sensing and motility. The ‘green’ module contained a single hub-like sORF_00203, which was associated with enrichment for carboxylic acid metabolism and lipid/terpenoid biosynthesis, with tRNA modification as a supporting process. Although unannotated, its consistent high MCC ranking and co-expression with metabolic genes suggest structural importance within the same metabolic framework.

While these network-based associations are informative, functional assignments here are based solely on GO-term enrichment and sequence homology and should be interpreted as predictive rather than confirmatory. The restricted hub-like set provides a more conservative and robust view of sORF network centrality, prioritizing candidates most likely to exert topological influence across modules. Direct biochemical characterization (for example, co-immunoprecipitation, ribosome profiling, or targeted proteomics) will be required to confirm the mechanistic roles of these hub sORFs.

## Discussion

In this study, we report a comprehensive genome-wide and transcriptomic analysis of small open reading frames (sORFs) in *Leptospira interrogans*, revealing their potential roles in regulatory, metabolic, and pathogenic processes in this species. By integrating sORF prediction, differential expression profiling, and co-expression network analysis, we uncover context-specific expression patterns and functional associations that suggest significant contribution of sORFs in the bacterium’s adaptability and virulence.

The high number of predicted sORFs in *L. interrogans* highlights an underexplored layer of the bacterial coding genome. Despite stringent filtering, most sORFs remain functionally unannotated, revealing significant gaps in current databases and annotation tools (Leong et al., 2022; Liu et al., 2022; Zapata et al., 2025). One important consideration is that sORFs can overlap with sRNAs. Although traditionally classified as noncoding, sRNAs have been shown to encode small peptides (Aoyama et al., 2022; George et al., 2024). By retaining sORFs overlapping with sRNAs, our analysis uncovered hub-like elements otherwise overlooked, such as sORF_00008 (‘blue’ module), which interacts with LIMLP-genes linked to motility and membrane remodeling, and energy processes.

Hub-like sORFs were identified based on intramodular topology rather than eigengene connectivity, using Maximal Clique Centrality (MCC) scores in CytoHubba. The top ten per module were designated as hub-like elements, reflecting their co-expression centrality rather than experimentally confirmed regulatory control. Only a restricted set of high-confidence hub-like candidates sORFs was retained, including sORF_0008 (‘blue’ module), sORF_00284 (‘purple’ module), sORF_00175 (‘red’ module), and sORF_00203 (‘green’ module), each occupying topological or central position indicative of network centrality rather than differential expression alone. Notably, in ‘plum1’, ‘purple’ and ‘skyblue’ modules, differentially expressed sORFs were recovered only when sRNA overlaps were considered. However, after applying the stricter MCC top-10 cutoff, these were not retained as hub-like nodes but remain functionally relevant for co-expression dynamics.

A total of 39 differentially expressed entirely novel sORFs, which could represent orphan genes. Such genes typically encode shorter proteins, approximately six-fold shorter than canonical proteins, with tissue- or condition-specific expression, simplified codon usage, and reduced amino acid complexity (Pereira et al., 2024). These sORFs display features consistent with lineage-specific or rapidly evolving gene families, including short length, reduced domain structure, and module-specific expression patterns. Younger orphan genes in other bacterial systems have been reported to evolve more rapidly than canonical genes, often contributing to lineage- or niche-specific adaptations (Vakirlis & Kupczok, 2024). Similar findings have been reported in other independent studies of Gram-negative bacteria, where orphan loci were shown to evolve under accelerated rates and to mediate lineage-specific traits such as host adaptation, environmental persistence, and pathogenicity (Ferrandis-Vila et al., 2022; Hussain et al., 2021). While we did not assess substitution rates or perform intraspecies genome comparisons in this study, these studies suggest that our identification of novel, differentially expressed sORFs in *L. interrogans* highlights putative candidates that may follow similar evolutionary trajectories. These findings underscore the value of integrative computational pipelines and emphasize the limitations of relying solely on sequence similarity for functional inference.

Transcriptomic profiling reinforces the functional relevance of sORFs in *L. interrogans*. The >32-fold upregulation of sORF_00332, despite its lack of functional annotation, underscores the limitation of sequence-based predictions and highlights the value of expression profiling for uncovering biologically significant elements. This transcriptional induction occurring in the *lomA* mutant background where virulence and motility are impaired as reported by Gaultney et al. (2020), suggesting that sORF_00332 may function in methylation-sensitive regulatory pathways or act as a master effector contributing to adaptive and pathogenic responses. Its identification highlights the potential of unannotated small proteins as high-priority candidates for experimental characterization, opening new avenues for integrating sORFs into bacterial regulatory networks directly relevant to virulence and translational applications.

Several other sORFs intersect with core cellular pathways including fatty acid metabolism, protein turnover, and signal transduction. For instance, sORF_00300 and sORF_00331, predicted to encode a Clp protease ATP-binding subunit protein and an adenosylhomocysteinase respectively, indicate the potential involvement of sORFs in proteostasis and methylation cycles that are both critical to stress response and bacterial adaptation (Aljghami et al., 2022; Koeppl et al., 2024). These functions align with the observed transcriptional shifts in canonical genes such as sigma-70 family RNA polymerase sigma factor and FecR domain-containing protein (LIMLP_RS03670 and LIMLP_RS03675, respectively), regulators of extracytoplasmic stress, which were upregulated in the mutant background (de Dios et al., 2021). Coordinated induction of iron acquisition genes, including energy transducer TonB and biopolymer transporter ExbD (LIMLP_RS04230 and LIMLP_RS04235, respectively), both components of TonB system further suggests that sORFs may participate in regulatory circuits that couple iron homeostasis with broader stress signaling (Grassmann et al., 2021). The ATP-binding motifs found in sORF_00166, hint towards the ABC transporter function protein family that is known for involvement in nutrient acquisition, antimicrobial resistance, and toxin secretion (Akhtar & Turner, 2022).

Upregulated DUF-domain-containing sORFs were observed to be linked to stress adaptation and host-pathogen interactions, suggesting that these sORFs could be involved in previously unrecognized virulence regulators. These findings support a role for sORFs as critical effectors of methylation-sensitive regulatory programs governing metabolic flexibility and virulence expression. The role of *lomA*-mediated 4mC DNA modification in *L. interrogans* was previously examined by Gaultney et al. (2020), who demonstrated that *lomA* inactivation leads to widespread transcriptional dysregulation, impaired growth, reduced cell adhesion, and complete loss of virulence in an acute infection model. Notably, they also reported diminished motility on semisolid medium but could not connect this phenotype to specific regulatory or structural pathways, as their analysis was limited to annotated differentially expressed genes. Building on this foundation, our study integrates sORFs into the transcriptome-wide regulatory landscape of *L. interrogans*. We confirm the broad transcriptional impact of *lomA* inactivation and identify differentially expressed sORFs that converge on motility-, virulence-, metabolic-associated pathways (Fig. 2). We propose that *LomA*-mediated methylation regulates a network of membrane-associated sORFs that control lipid remodeling and flagellar anchoring, providing a mechanistic hypothesis for the motility defects previously observed (Fig. 3).

**Fig. 3.**
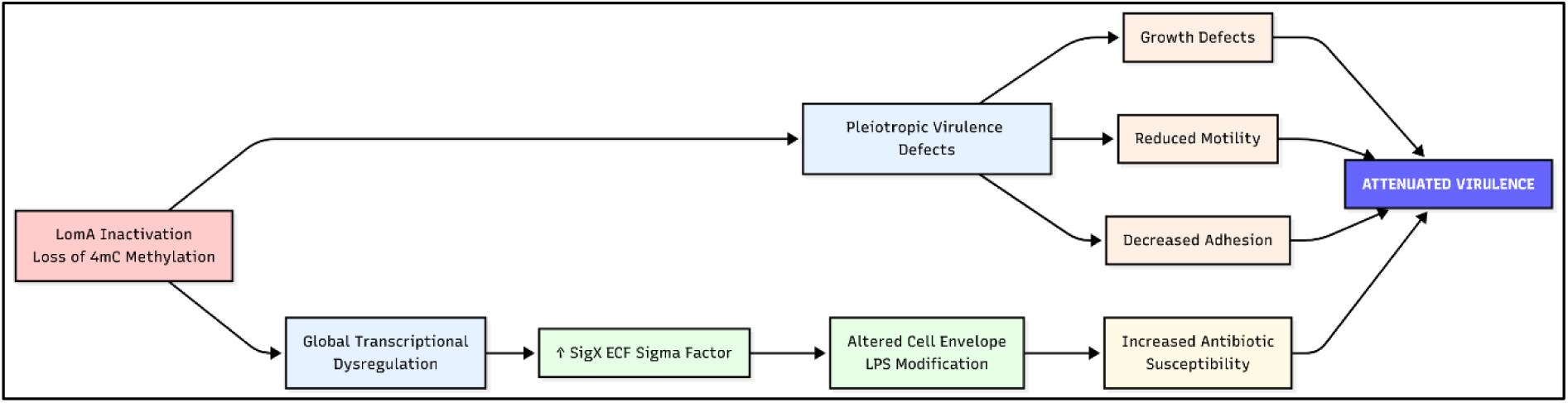
Proposed model of *LomA*-dependent 4mC methylation and its contribution to global regulation and virulence in *L. interrogans*. Loss of *LomA*-mediated 4mC methylation results in transcriptional dysregulation, including induction of the extracytoplasmic function sigma factor SigX and changes in cell envelope and LPS-modification pathways. These effects lead to growth defects, reduced motility, decreased adhesion, and increased antibiotic susceptibility, culminating in attenuated virulence. This model synthesizes previous phenotypic findings with the transcriptomic and sORF-based regulatory insights generated in this study.

Motility and chemotaxis emerged as dominant functional themes across modules, underscoring their centrality in the phenotypic outcomes of *lomA* disruption. In the ‘blue’ module, upregulation of sORF_00022 (DUF-transport protein) and sORF_00128 (GtrA glycosyltransferase), together with the surface SBBP repeat-containing protein (LIMLP_RS02105), formed networks enriched for flagellar assembly and basal body organization. Their predicted interactions with the putative hub-like sORF_00008 (lysophospholipid acyltransferase) suggest a lipid-remodeling mechanism that secures flagellar insertion and stability, consistent with analogous processes in *Pseudomonas* and *Vibrio* spp. (Deryusheva et al., 2023; Fulton et al., 2024; Homma & Kojima, 2022; Sánchez-Peña et al., 2024). Consistent with this, the retained hub-like sORFs (particularly sORF_00008, sORF_00175, and sORF_00203) represent nodes bridging motility, environmental sensing, and metabolic adaptation rather than broad module-wide regulatory centers.

Chemotaxis-related processes were also represented across multiple modules. In the ‘red’ module, sORF_00175 co-expressed with cheR and an M23 metallopeptidase, linking envelope remodeling to chemotactic signaling. Its interaction with an ABC-F protein suggests a role in maintaining translation of motility-associated proteins under stress, a mechanism reported in Gram-negative pathogens (Fostier et al., 2021). Similarly, sORF_00203 in the ‘green’ module connected lipid and carboxylic acid metabolism with tRNA modification, suggesting a potential link between metabolic adaptation and chemotactic function. These patterns reinforce a systems-level view in which sORFs align chemotaxis with envelope integrity and metabolic adaptation. Consistent with MCC-based ranking, sORF_00203 was the single retained hub-like sORF within its module, associated with carboxylic acid metabolism and lipid/terpenoid biosynthesis. Its co-expression with tRNA modification genes implies functional coupling between energy metabolism and translational regulation. In contrast, hub-like sORF_00284 (‘purple’ module) was associated with methyltransferase activity and nucleotide biosynthetic processes, aligning with roles in metabolic regulation and potential epigenetic modification.

Earlier analyses yielded a broader hub-like set, but the refined MCC-based restriction to the top ten nodes per module provides a more conservative yet biologically coherent framework. These hub-like sORFs (sORF_00008, sORF_00284, sORF_00175, and sORF_00203) represent robust network anchors linking motility, chemotaxis, and metabolism under *LomA*-dependent regulation. This refinement enhances confidence in topological inference and avoids overinterpretation of co-expression connectivity. Taken together, this study provides a genome-wide systems perspective on sORFs in *L. interrogans*, demonstrating their integration into motility, chemotaxis, genome stability, energy metabolism, and envelope biogenesis. By incorporating unannotated coding elements into network-based analyses, we contextualize the reduced motility phenotype reported by Gautney et al. (2021), linking it to structural and regulatory circuits involving lipid remodeling, glycosylation, and energy metabolism. The refined hub-like set provides a high-confidence framework for future validation, highlighting key sORFs most likely to exert regulatory influence, in contrast to analyses based solely on differential expression.

The identification of numerous differentially expressed orphan candidates underscores major gaps in bacterial genome annotation and supports the emerging view that small proteins can serve as critical regulatory and structural elements. While these insights are based on computational inference, targeted genetic, transcriptomic, and proteomic validation will be essential to confirm their mechanistic roles. Such sORFs could provide novel targets for diagnostic or therapeutic intervention, with potential to disrupt key adaptive and pathogenic pathways in leptospiral infection. Beyond deepening our understanding of bacterial regulatory biology, these findings hold translational relevance for mitigating the public health and economic impact of leptospirosis.

## Conclusion

In summary, our genome-wide and transcriptomic analyses reveal that sORFs in *L. interrogans* occupy central regulatory positions in co-expression networks and respond dynamically to epigenetic perturbation. The identification of hub-like sORFs linking motility, chemotaxis, metabolism, and membrane remodeling provides candidate targets for functional interrogation. Differentially expressed orphan sORFs further highlight previously unrecognized regulatory elements that may contribute to pathogen adaptability and virulence. From a translational perspective, these microproteins present opportunities for the development of novel antimicrobial strategies, including targeted peptide or small-molecule inhibitors, antisense therapeutics, and epitope-based vaccines aimed at disrupting critical bacterial adaptive circuits. Beyond therapeutic applications, the insights gained into sORF-mediated regulatory architecture may inform synthetic biology approaches for designing robust microbial chassis or biosensors. Collectively, this work not only expands fundamental understanding of bacterial microproteins but also underscores their potential impact on public health and disease mitigation strategies for leptospirosis.

## Acknowledgement

We acknowledge the contributions of all authors to this work. Chung Yuen Khew and Dr. Norfarhan Mohd-Assaad led the analysis and manuscript preparation. Nur Syafiqah Mohd Fowzi, Nurul Najihah Zaifulzaman and Chung Yuen Khew performed the data reanalysis and contributed to the interpretation of results. Dr. Sarahani Harun, Dr. Shairah Abdul Razak and Prof. Dr. Zeti-Azura Mohamed-Hussein provided critical feedback, proofreading and conceptual input that improved the clarity and rigor of the manuscript. Dr. Norfarhan Mohd-Assaad supervised the study and served as the corresponding author. We thank all contributors for their effort and collaboration.

## Statements and Declarations

### Author Contributions

All authors contributed to the study conception and design. Material preparation, data collection and analysis were performed by ChungYuen Khew, Nur Syafiqah Mohd Fowzi, and Nurul Najihah Zaifulzaman. The first draft of the manuscript was written by ChungYuen Khew, and all authors commented on previous versions of the manuscript. Dr. Sarahani Harun, Dr. Shairah Abdul Razak, and Prof. Dr. Zeti-Azura Mohamed-Hussein provided critical review, proofreading, and conceptual input that strengthened the manuscript. Dr. Norfarhan Mohd-Assaad supervised the study and served as the corresponding author. All authors read and approved the final manuscript.

### Data Availability

The datasets analyzed in this study’s findings are available in the Gene Expression Omnibus genomics data repository, under GEO accession number GSE138917.

### Ethics approval

This study did not involve new experiments with human participants, animals, or recombinant DNA. All analyses were performed on previously published and publicly available sequencing data. Ethical approval was therefore not required.

### Consent to participate

Not applicable. The study did not involve human participants.

### Consent to publish

Not applicable. The study did not involve human participants and uses only previously published sequencing datasets.

